# *Allicin* regulates energy homeostasis through brown adipose tissue

**DOI:** 10.1101/713305

**Authors:** Chuanhai Zhang, Xiaoyun He, Yao Sheng, Jia Xu, Cui Yang, Shujuan Zheng, Junyu Liu, Haoyu Li, Jianbing Ge, Minglan Yang, Baiqiang Zhai, Wentao Xu, Yunbo Luo, Kunlun Huang

**Affiliations:** Beijing Advanced Innovation Center for Food Nutrition and Human Health, College of Food Science and Nutritional Engineering, China Agricultural University, Beijing, 100083, China; Key Laboratory of Safety Assessment of Genetically Modified Organism (Food Safety), Ministry of Agriculture, Beijing, 100083, China

**Keywords:** Allicin, energy homeostasis, brown adipose tissue, obesity, mitochondria

## Abstract

**Background/objectives:** Disorder of energy homeostasis can lead to a variety of metabolic diseases, especially obesity. Brown adipose tissue (BAT) is a promising potential therapeutic target for the treatment of obesity and related metabolic diseases. *Allicin*, a main bioactive ingredient in garlic, has multiple biology and pharmacological function. However, the role of *Allicin*, in the regulation of metabolic organ, especially the role of activation of BAT, has not been well studied. Here, we analyzed the role of Allicin in whole-body metabolism and the activation of BAT.

**Results:** Allicin had a significant effect in inhibiting body weight gain, decreasing adiposity, maintaining glucose homeostasis, improving insulin resistance, and ameliorating hepatic steatosis in diet-introduced obesity (DIO) mice. Then we find that Allicin can strongly activate brown adipose tissue (BAT). The activation of brown adipocyte treated with Allicin was also confirmed in mouse primary brown adipocytes.

**Conclusion:** Allicin can ameliorate obesity through activating brown adipose tissue. Our findings provide a promising therapeutic approach for the treatment of obesity and metabolic disorders.

## Introduction

The regulation of energy homeostasis, including food intake, energy expenditure, and body adiposity, is crucial for the health of the body (*1, 2*). Obesity is a result of unbalance of energy homeostasis which caused by energy intake exceeds energy expenditure, and then excess energy stored in fat in the form of triglycerides which lead to adiposity(*3*) and can cause a series of metabolic diseases, such as insulin resistance, type 2 diabetes mellitus cardiovascular disease, cancer, inflammation and other related diseases(*4*). Currently, obesity has become one of the fastest-growing non-communicable diseases, and it estimated that by 2030, 38% of adults would be overweight and 20% will develop obesity in the world (*5, 6*). Although the situation of obesity is dire, satisfactory safe and effective treatment strategies are limited for individuals(*7*).

There are two main types of fat in mammals, including energy-storing white adipose tissues (WAT) and energy-consuming brown adipose tissues (BAT) (*8*). Different from WAT, BAT contains a large number of mitochondria which is responsible for nonshivering thermogenesis in mammals(*9, 10*). Increasing studies showed that the potential therapy of anti-obesity and related metabolic diseases are related to the activation of BAT, which bring a new way to fight obesity(*11–14*). Besides, the proper stimulus could induce the generation of UCP1 (uncoupling protein-1) in WAT, which is also rich in mitochondria and defined as beiging (*15*). Many studies suggested that beiging in WAT also can effectively enhance energy expenditure and improve metabolic disorder(*16, 17*). Furthermore, reducing the amount of WAT and the size of white adipocyte can effectively improve inflammation and metabolic disorders (*18*). Both WAT and BAT have essential functions in regulating energy homeostasis. Therefore, increasing activation of BAT and induction of *beiging* in WAT could be an effective therapeutic strategy for obesity and its related diseases.

*Allicin*, a main bioactive ingredient in garlic(*19*), has multiple pharmacological functions, including anti-oxidative stress, anti-tumor, cholesterol-lowering, anti-platelet aggregation, liver protection, prevention of cardiovascular disease and anti-inflammatory (*20–22*). Although Allicin induces beiging in WAT(*23*), the effect of the regulation of energy homeostasis including whole-body energy metabolism and the mechanism of activation of BAT of *Allicin* is still unclear.

In this study, we determined the effect of the whole-body energy metabolism and activating BAT of Allicin.Our data establish that Allicin plays a previously not fully recognized role in regulating energy homeostasis, which may provide more rigorous thinking of the therapeutic approach to the treatment of metabolic disorders.

## 2. Materials and methods

### 2.1. Animals

C57BLKS/J-Leprdb/Leprdb (Db/Db) male mice (3-weeks old) purchased from the Model Animal Research Center of Nanjing University. Male C57BL/6J mice (15–20 g, 3-4 weeks old) were purchased from Vital River Laboratories (Beijing, China) and housed under 22±2°C and 55%±10% humidity with 12-hour light-dark cycle.

For the DIO mice model, eight mice were fed a low-fat diet (LFD), and the rest mice were fed high-fat-diet (HFD) until the weight difference was over 18-20% for six weeks (Figure s1A). We marked the mice fed an LFD and sterile water as the vehicle control group (LFD, n = 8). The HFD mice divided into four groups: control group (HFD, n = 8) fed an HFD and sterile water, and three treatment groups fed an HFD and ALLICIN solutions (Allic+High, Allic+Mid, Allic+Low with 0.3%, 0.6%, 1% m/m ALLICIN solution for drinking, respectively, n = 8). The experiment lasted for another 13 weeks. ALLICIN solutions refreshed every 1-2 days. ALLICIN (98%, YZ-100384, Solarbio) purchased from Solarbio Life Science Biotechnology Co. Ltd. All experimental procedures conducted and the animals used according to the Guide for the Care and Use of Laboratory Animals published by the U.S. National Institute of Health and approved by the Animal Ethics Committee of China Agricultural University, Beijing.

### 2.2. Glucose tolerance test and insulin tolerance test

GTT and ITT performed during the last week of the experiment. For GTT, mice fasted for 16h with free access to drinking water. Glucose (0.8 g/kg for Db/Db mice and 1.5 g/kg for HFD and LFD mice) was administered intraperitoneally, and blood glucose levels were measured with an Accu-Chek glucose monitor (ACCU-CHEK, Shanghai, China) at 0, 15, 30, 60, 90 and 120 min. For the insulin tolerance test (ITT), the mice fasted for 4 h (9:00 AM to 1:00 PM), and the mice were administered human insulin (0.7 U/kg Humulin R; Novo Nordisk) by intraperitoneal injection. Blood glucose concentrations were determined from the tail vein with a blood glucose meter (ACCU-CHEK) at 0, 15, 30, 45, and 60 min after the insulin injection.

### 2.3. Cold-induced thermogenesis

For cold tolerance test, mice were placed in a cold chamber (4°C) for four h. We evaluated the cold-induced thermogenesis by measuring the rectal temperature with a temperature sensor (AT210, Zhongyidapeng, Shenzhen, China). Then we took infrared thermal imaging of the animals by a handheld infrared camera (FLIR T600) on a whiteboard.

### 2.4. Metabolic rate and physical activity

Oxygen consumption and physical activity were determined at the 12th week of the experiment with a TSE lab master (TSE Systems, Germany) (*24*). Mice were acclimated to the system for 20–24h, and then we measured oxygen consumption (VO_2_ and VCO_2_) over the next 24 h. The animals were maintained at 25°C in a 12-h light/dark cycle with free access to food and water. We measured the voluntary activity of each mouse using an optical beam technique (Opto-M3, Columbus Instruments, Columbus, OH, USA) over 24 h and expressed as 24 h average activity. The respiratory exchange ratio (RER) was then calculated (*25*).

### 2.5. Body composition measurements

The total fat and lean masses of mice after a 12-week treatment with either vehicle or ALLICIN were assessed with the Small Animal Body Composition Analysis and Imaging System (MesoQMR23 - 060H-I, Nuimag Corp., Shanghai, China), according to the manufacturer’s instructions.

### 2.6. Positron emission-computed tomographic imaging

Specific method reference from Yuan et al. Briefly described as the mice were left unfed overnight and was lightly anesthetized with isoflurane followed by a tail vein injection of fluorodeoxyglucose ([18F]-FDG; 500 mCi). They were subjected to PET-CT analysis at 60min after radiotracer injection. Inveon Acquisition Workplace (IAW) software (Siemens Preclinical Solutions) was used for the scanning process. A 10-min CTX-ray for attenuation correction was scanned with a power of 80 kV and 500 mA and an exposure time of 1100ms before the PET scan. Ten-minute static PET scans were acquired, and images were reconstructed by ordered-subsets expectation maximization (OSEM) 3-dimensional algorithm followed by maximization/maximum a posteriori (MAP) or Fast MAP provided by IAW. The 3-dimensional regions of interest (ROIs)were drawn over the guided CT images, and the tracer uptake was measured with the IRW software (Siemens). Specific quantification of the [18F]-FDG uptake in each of the ROIs was calculated. The data for the accumulation of [18F]-FDG on micro-PET images were expressed as the standard uptake values, which were determined by dividing the relevant ROI concentration by the ratio of the injected activity to the bodyweight.

### 2.7. Histology

Tissues fixed with 4% paraformaldehyde were sectioned after embedment in paraffin. We prepared multiple sections for hematoxylin-eosin staining. We incubated cells grown on poly-L-lysine-pretreated coverslips (Sigma-Aldrich, St. Louis, MO, USA) in 5% goat serum for 1 h at room temperature after fixation with 1% formalin at room temperature for 1 h. All images were acquired with the BX51 system (Olympus, Tokyo, Japan).

### 2.8. Brown adipocyte differentiation

The isolation method of mouse BAT primary cells referred to(*26*). Primary cells were cultured in basal medium (containing 80% DMEM, 20% fetal bovine serum, and 1% penicillin and streptomycin) until the cells had reached more than 90% confluency. Then, cells were treated with a brown adipogenic induction cocktail (DMEM) containing 10% fetal bovine serum, 1% penicillin and streptomycin, 20 nM insulin, 1 mM dexamethasone, 0.5 mM isobutylmethylxanthine, 125 nM indomethacin, and 1 nM 3,3,5-triiodo-L-thyronine (T3) for the first two days. The medium was then replaced by the medium supplemented with only insulin and T3 every other day. The cells were treated with or without ALLICIN (50 μg/mL, SA8720, Solarbio) for six days during brown adipogenesis. BAT differentiation medium was used as the solvent, and 0.1 μg/mL, 1 μg/mL, 10 μg/mL, 50 μg/mL and 100 μg/mL solutions were used as the material for the *in vitro* tests. At day 6, fully differentiated adipocytes were used for the experiments.

### 2.9. Measurements of cellular respiration

BAT primary adipocytes were treated with and/or without Allicin (50 μg/mL) at day 6 of brown adipogenesis for 24h. We measured O_2_ consumption of fully differentiated adipocytes at day 7 with an XF24-3 extracellular flux analyzer (Agilent Technologies, Santa Clara, CA, USA). Basal respiration was also assessed in untreated cells.

### 2.10. RNA isolation and real-time quantitative PCR

Total RNAs from tissues or cells were extracted with Trizol reagent (Thermo Fisher Scientific). We used equal amounts of RNA to synthesize cDNA with the transcript One-Step gDNA Removal and cDNA Synthesis SuperMix kit (AT311-03, TransGen Biotech, Beijing, China). The PCR reaction was run in triplicate for each sample using a Prism VIIA7 real-time PCR system (Thermo Fisher Scientific). Primer sequences are available upon request.

### 2.11. Western blot analysis

An equal amount of protein from cell lysate was loaded into each well of a 12%SDS-polyacrylamide gel after denaturation with SDS loading buffer. After electrophoresis, we transferred proteins to a PVDF membrane and incubated with blocking buffer (5% fat-free milk) for 1 h at room temperature. The following antibodies were added overnight: anti-UCP1 (Abcam, ab10983, 1:1000 diluted in 5% BSA, 0.0025% Tween-20 in 1× TBS solution), anti-Oxphos (Abcam, ab110413, 1:1000 dilution), anti-β-actin (CST, #8457, 1:1000 dilution) and anti-β-tubulin (CST, #2146, 1:1000 dilution). These primary antibodies were incubated overnight in a 4°C refrigerator. The membrane was incubated with horseradish peroxidase-conjugated secondary antibodies for 1 h at room temperature. All signals were visualized and analyzed by Clinx Chemi Capture software (Clinx, Shanghai, China).

### 2.12. Immunofluorescence staining

We stained differentiated cells with anti-human UCP1, followed by an Alexa 488-conjugated secondary antibody (Invitrogen, Carlsbad, CA, USA), BODIPY (Thermo Fischer Scientific, Waltham, MA, USA) and DAPI (Leagene, Beijing, China), complying with manufacturers’ the procedure. Brown adipocytes were positive for both UCP1 and BODIPY. We stained negative controls with the omission of a primary antibody. Images were taken by Zeiss laser scanning confocal microscopy (LSM780, Oberkochen, Germany).

### 2.13. Statistical analysis

We used a single-factor analysis of variance (ANOVA) followed by a two-tailed Student’s t-test for comparisons. We presented almost all data as means ± SEM. Significant differences were considered when *P* < 0.05.

## Result

### 3.1. Allicin reduces adiposity and maintains glucose homeostasis in mice

To investigate the effect of allicin on energy homeostasis, the high-fat-diet-induced obesity mice model (HFD) and genetically leptin-receptor-deficiency obese (Db/Db) mice model was used to assess the role of Allicin on obesity and energy metabolism. Firstly, we screened the optimal treatment concentration (Allicin-high) of allicin by observing the effect of three doses of allicin on the reduction of body weight of HFD mice *(Fig. 1A and Fig. s1B)* and also treated the Db/Db mice with the optimal concentration. A significant reduction in body weight after Allicin treatment was observed both in HFD and Db/Db mice models *(Fig. 1A–1B, Fig. 2A–2C)*. Notably, the protection from weight gain after Allicin treatment mainly due to the reduction of fat accumulation in eWAT and sWAT *(Fig. 1C–1E, Fig. 2D and 2E)*. These data indicated that the anti-adiposity effect of Allicin is evident from both long-term accumulations of innate and genetic obesity. Besides, adiposity often affects glucose homeostasis, so glucose homeostasis and insulin resistance were analyzed by glucose tolerance testing (GTT) and insulin tolerance testing (ITT) both in HFD and Db/Db mice. As expected, Allicin treatment improved glucose tolerance and insulin sensitivity *(Fig. 1F–1G, Fig. 2F–2G)*. Correspondingly, Serum profiles, including total cholesterol (CHO), TG, LDL, and NEFA levels, were also markedly reduced after Allicin treatment *(Fig. s4)*. Taken together, these results indicated that Allicin notably ameliorates adiposity and maintains glucose homeostasis in mice.

**Figure 1.**
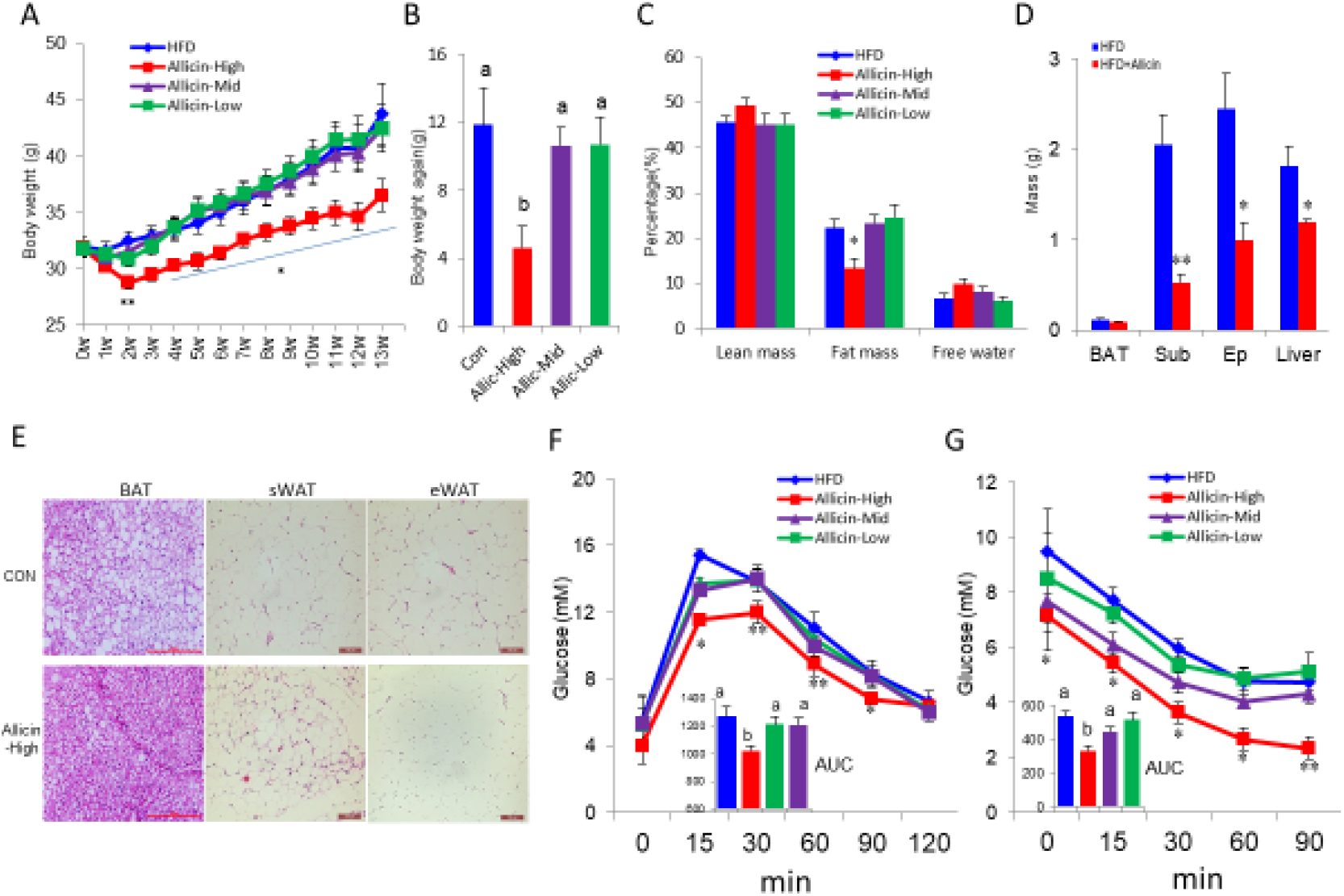
Allicin reduces adiposity and improves glucose homeostasis in DIO mice. Vehicle (control) or different-doses Allicin were daily administrated for 13 weeks. (A) Bodyweight evaluation of control HFD mice or mice treated with different-dose Allicin. (n=9). (B) Bodyweight again of different groups of mice (n=9). (C) Body fat percentage test using NMR. (D) Organ weight of control and Allicin-treated HFD mice (n = 6). (E) H&E staining of BAT, sWAT, and eWAT sections from DIO control and Allicin treated DIO mice. (F) Glucose tolerance test on control and Allicin-treated DIO mice (I.P. with glucose as 1.0 g/kg after 16h fast) (n=8). Also, the average area under the curve (n = 8). (G) Insulin tolerance test was performed on control and Allicin-treated DIO mice (injection insulin with 2.0 U/kg after 6 h of fasting (n = 8). And the average area under the curve (n=8). Values represent means ± SEM. Error bars represent SEM; significant differences compared to vehicle controls are indicated by *p < 0.05, **p < 0.01, ***p < 0.001 (assessed by Student’s t-test).

**Figure 2.**
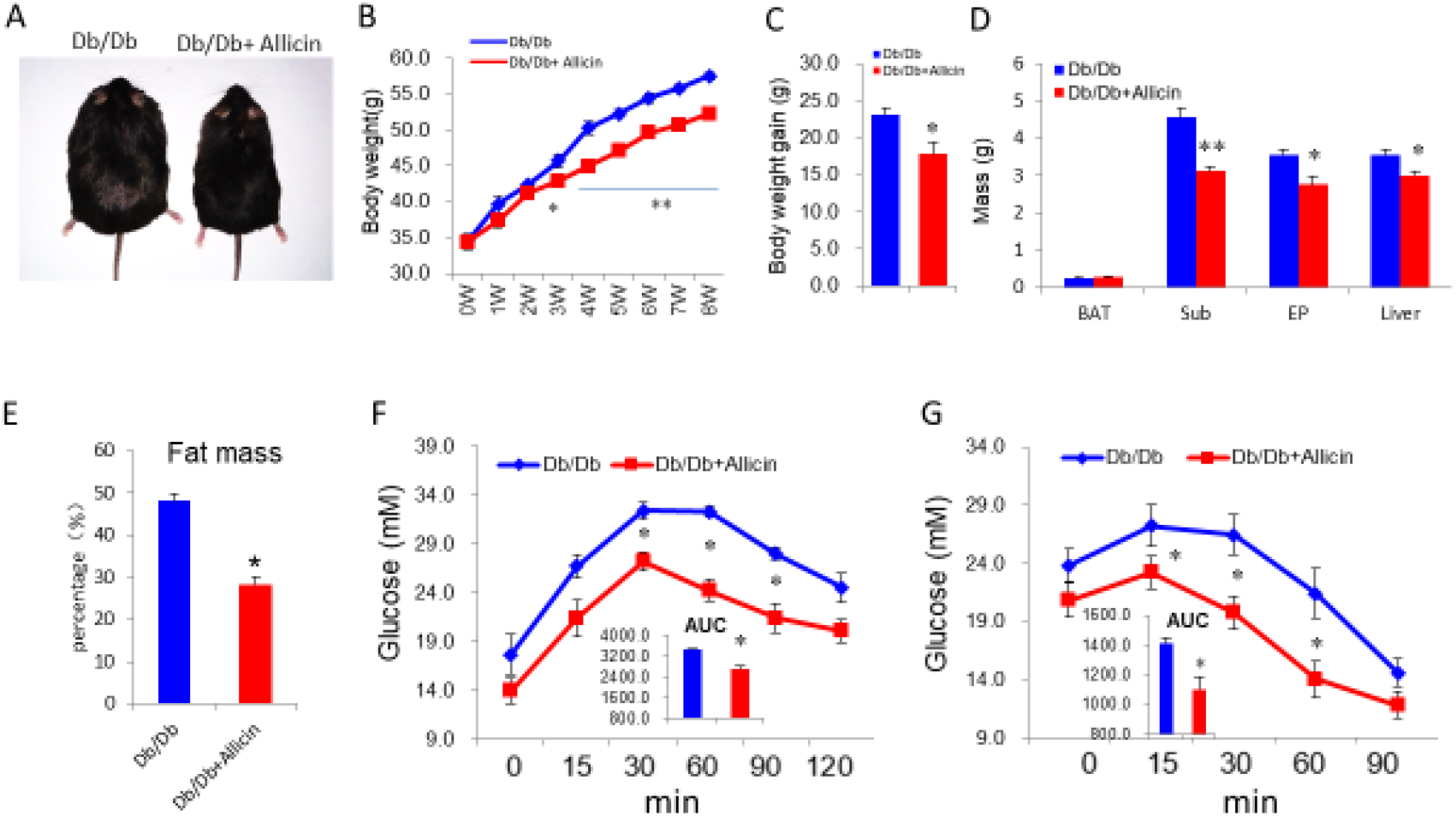
Allicin reduces body weight gain and improves glucose homeostasis in leptin-receptor deficiency mice (Db/Db mice). (A) Body-shape image of Db/Db control mice and mice treated with Allicin. (B) Bodyweight of Db/Db mice treated with vehicle or Allicin. (n=5). (C) Bodyweight gain of Db/Db control mice and mice treated with Allicin. (n=5). (D) Organ weight of control and Allicin-treated Db/Db mice (n = 5). (E) Body fat percentage test using NMR. (F) Glucose tolerance test on control and Allicin-treated Db/Db mice (I.P. with glucose as 0.5 g/kg after 16h fast) (n=5). And the average area under the curve (n = 5). (G) Insulin tolerance test was performed on control and Allicin-treated Db/Db mice (injection insulin with 1.0 U/kg after 6 h of fasting (n = 5). And the average area under the curve (n=5). Values represent means ± SEM. Error bars represent SEM; significant differences compared to vehicle controls are indicated by *p < 0.05, **p < 0.01, ***p < 0.001 (assessed by Student’s t-test).

### 3.2. Allicin augments whole-body energy expenditure by activation of brown adipose tissue

Energy consumption is an essential indicator of assessing energy homeostasis. We next investigated the effect of Allicin on whole-body energy expenditure in HFD and Db/Db mice with *the respiratory metabolic system*. We found that Allicin markedly increased the oxygen consumption both in HFD and Db/Db mice *(Fig. 3A–3C, Fig. 4A and 4B)*, indicating that Allicin-treated group mice have a higher energy expenditure. However, there is no significant difference in physical activity, food intake, or water intake *(Fig. 3D–3E, Fig. S2)*. Besides, Allicin treatment significantly increased the thermogenesis of interscapular BAT site and rectal temperature after cold stimulation in all Allicin treatment groups compared with the control group both in HFD and Db/Db mice *(Fig. 3G and 3H, Fig. 4C and 4D*). BAT is well known as an essential thermogenic and energy-consuming organ in the body, maintaining the body temperature by non-shaking thermogenesis under the stimuli of cold. Based on the above results, we hypothesized that Allicin augments whole-body energy expenditure by activation of BAT in HFD and Db/Db mice. Expectedly, Allicin treatment significantly induced the expression of genes related to thermogenesis and energy expenditure, including UCP1, PRDM16, PGC1α/β, CPT1α, and MCAD, in the BAT from HFD mice *(Fig. 3I)*.

**Figure 3.**
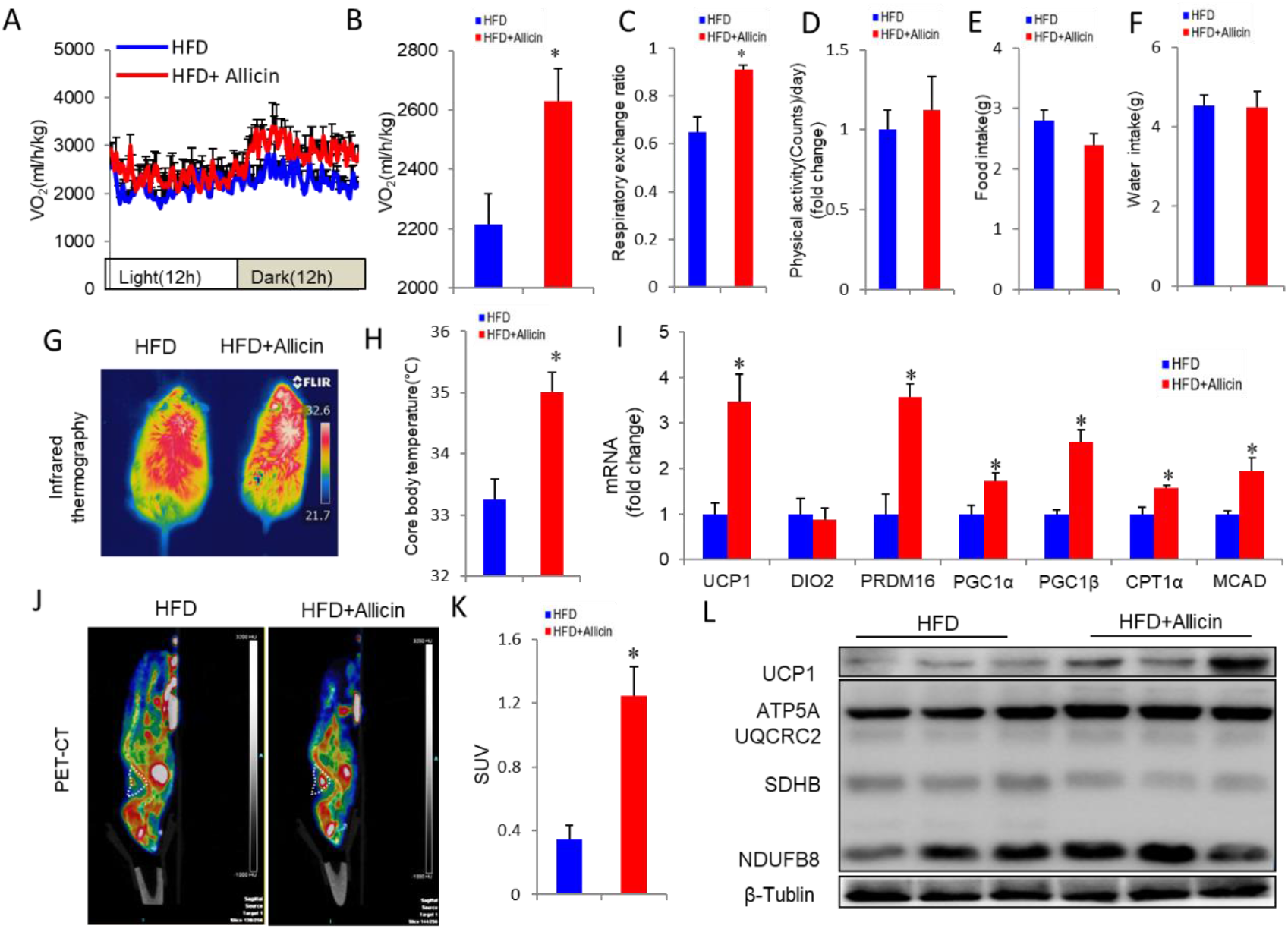
Allicin increases energy expenditure and enhances BAT activity in DIO mice. (A and B) Energy expenditure was assessed by oxygen consumption (VO2) in DIO mice after 12 wk of Allicin treatment (B) (n = 5); (C-F) physical activity (C); respiratory exchange ratio (D); food intake (E); water intake (F) of control and Allicin-treated DIO mice (n = 5). (G) Infrared thermal images of control and Allicin-treated DIO mice, which showed more thermal signals in the interscapular BAT position. (H) The core body temperature of control and Allicin treated mice after cold stimulation (4°C for 4 h). (I) The real-time PCR analysis of thermogenic genes in BAT form control and Allicin-treated DIO mice. (J-K) PET/CT images after injection of 18F-FDG into DIO mice treated with vehicle and Allicin for 12 weeks (J).white triangle indicates the anatomical site of the interscapular BAT (n=3). Mean standard uptake value (SUV) of 18F-FDG in BAT (K). UCP1 and OXPHOS expression levels in BAT from control and Allicin-treated DIO mice (L). Values represent means ± SEM. Error bars represent SEM; significant differences compared to vehicle controls are indicated by *p < 0.05 (assessed by Student’s t-test).

**Figure 4.**
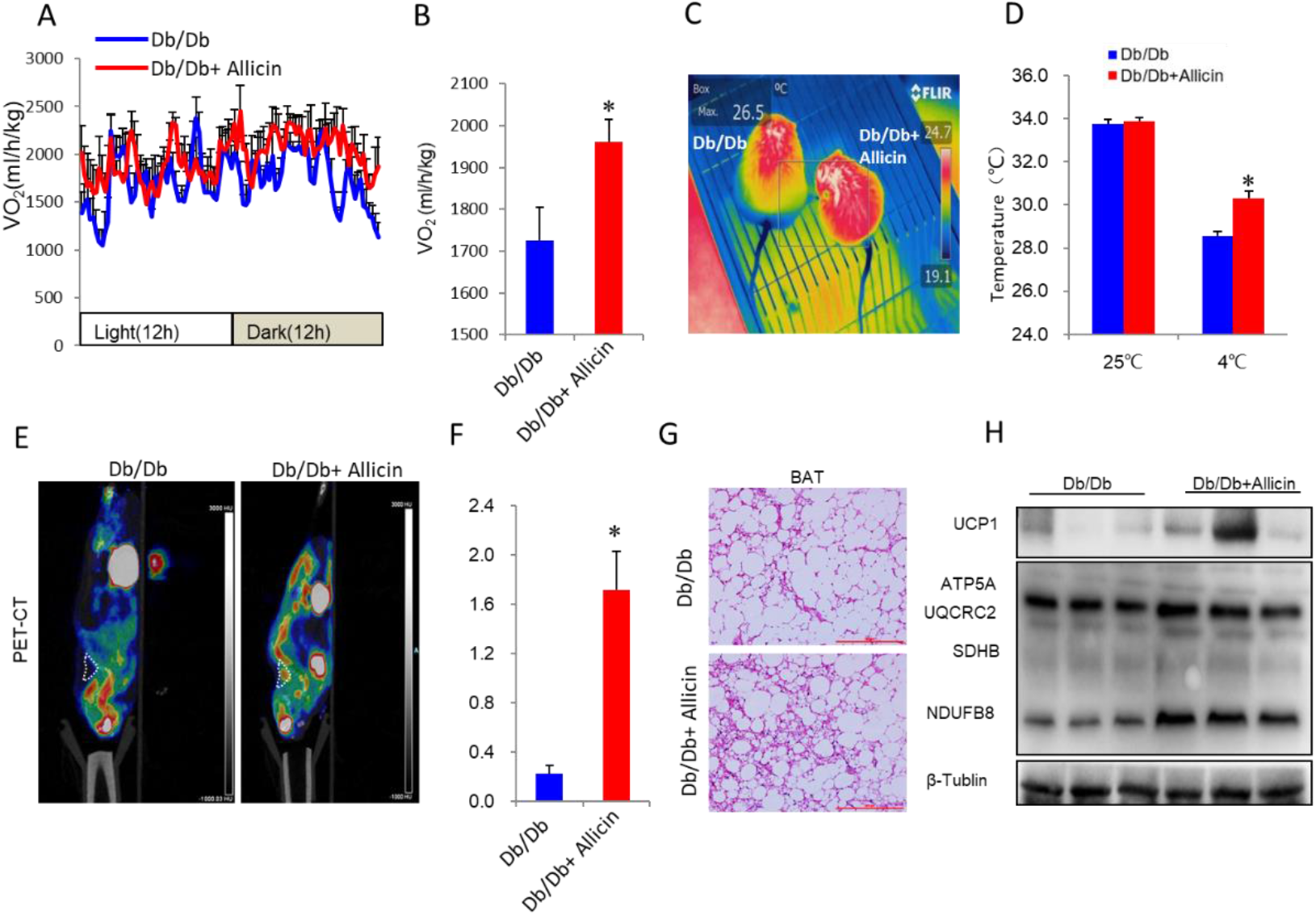
Allicin increases energy expenditure and enhances BAT activity in leptin-receptor deficiency mice (Db/Db mice). (A and B) Energy expenditure was assessed by oxygen consumption (VO2) in Db/Db mice after 8 weeks of Allicin treatment (B) (n = 5); (C) Infrared thermal images of control and Allicin-treated Db/Db mice, which showed more thermal signals in the interscapular BAT position. (D) The core body temperature of control and Allicin treated mice at room temperature (25°C) and after cold stimulation (4°C for 4 h). (E-F) PET/CT images after injection of 18F-FDG into Db/Db mice treated with vehicle and Allicin for 8weeks (E).white triangle indicates the anatomical site of the interscapular BAT (n=3). Mean standard uptake value (SUV) of 18F-FDG in BAT (F). (G) H&E staining of BAT form control and Allicin-treated Db/Db mice. (H) UCP1 and OXPHOS expression levels in BAT from control and Allicin-treated Db/Db mice. Values represent means ± SEM. Error bars represent SEM; significant differences compared to vehicle controls are indicated by *p < 0.05 (assessed by Student’s t-test).

Meanwhile, the 18F-fluorodeoxyglucose(FDG) PET combined with X-ray CT analysis shows that Allicin treatment dramatically increased the glucose utilization rate both in HFD and Db/Db mice *(Fig.. 3J and 3K, Fig. 4E and 4F)*. Histological analysis of BAT indicates that allicin treatment can reduce the size of lipid droplet *(Fig. 2F and Fig. 4G)*. The immunoblotting analysis further indicated that Allicin significantly enhanced the UCP1 and OXPHPS-related proteins (ATP5A, UQCRC2, SDHB, and NDUFB8) expression in BAT both from HFD and Db/Db mouse *(Fig. 3L and Fig. 4H)*. These results strongly indicated that that Allicin dramatically increased BAT activity in the HFD and Db/Db mice. These results altogether confirm that Allicin treatment increased *whole-body energy expenditure* and reduced adiposity in the mice.

### 3.3. Allicin active the brown adipocytes and increase the energy expenditure in vitro

The above results have proved that Allicin can increase energy expenditure through the activation of BAT in vivo. However, it is unclear whether allicin activates brown fat as a direct or indirect effect. In order to Fig. out this question, we isolated brown primary adipocytes from mice and performed an in vitro brown adipocytes differentiation assay with and/or without Allicin treatment. As the results, Allicin increased the expression of UCP1, the golden maker of activation of BAT, in a dose-dependent manner and with best activation effect at a concentration of 50 μM *(Fig. 5A)*. Consistently, the thermogenic genes, including PGC1α/β and CPT1α/β were also upregulated after the Allicin treatment compared with the solvent treatment *(Fig. 5B)*. Besides, UCP1 levels were further confirmed by quantification of the protein expression by immunoblotting and immunostaining *(Fig. 5C and 5D)*. Furthermore, the expression levels of OXPHOS proteins, including ATP5A, UQCRC2, SDHB, and NDUFB8, were markedly upregulated after Allicin treatment compared with the solvent treatment *(Fig. 5D)*. Importantly, the effect of Allicin on energy expenditure were accessed by an oxygen consumption rate (OCR) in brown adipocytes. Expectedly, the OCR-related basal metabolic rate, ATP levels, maximum oxygen consumption, and proton leakage were all markedly increased after Allicin treatment compared with solvent treatment *(Fig. 5E–4I)*. Notably, we also investigated the effect of Allicin on activation of brown adipocytes in human brown adipocytes. Surprisingly, the expression levels of thermogenic genes including UCP1, PGC1α/β, and CPT1α/β were dramatically increased after the brown adipocyte differentiation cocktail medium treatment with Allicin compared with solvent *(Fig. s5A)*. Correspondingly, the UCP1 levels in human brown adipocytes were further confirmed by immunoblotting and immunostaining *(Fig. s5D and s5E)*. Meanwhile, the expression levels of OXPHOS proteins in human brown adipocytes also were significantly upregulated after Allicin treatment compared with the solvent treatment *(Fig. s5E)*. The human brown adipocytes results indicated that Allicin also has the potential to be applied to human metabolic disease. Collectively, these results indicate that Allicin can active brown adipocytes and increase the energy expenditure in vitro.

**Figure 5.**
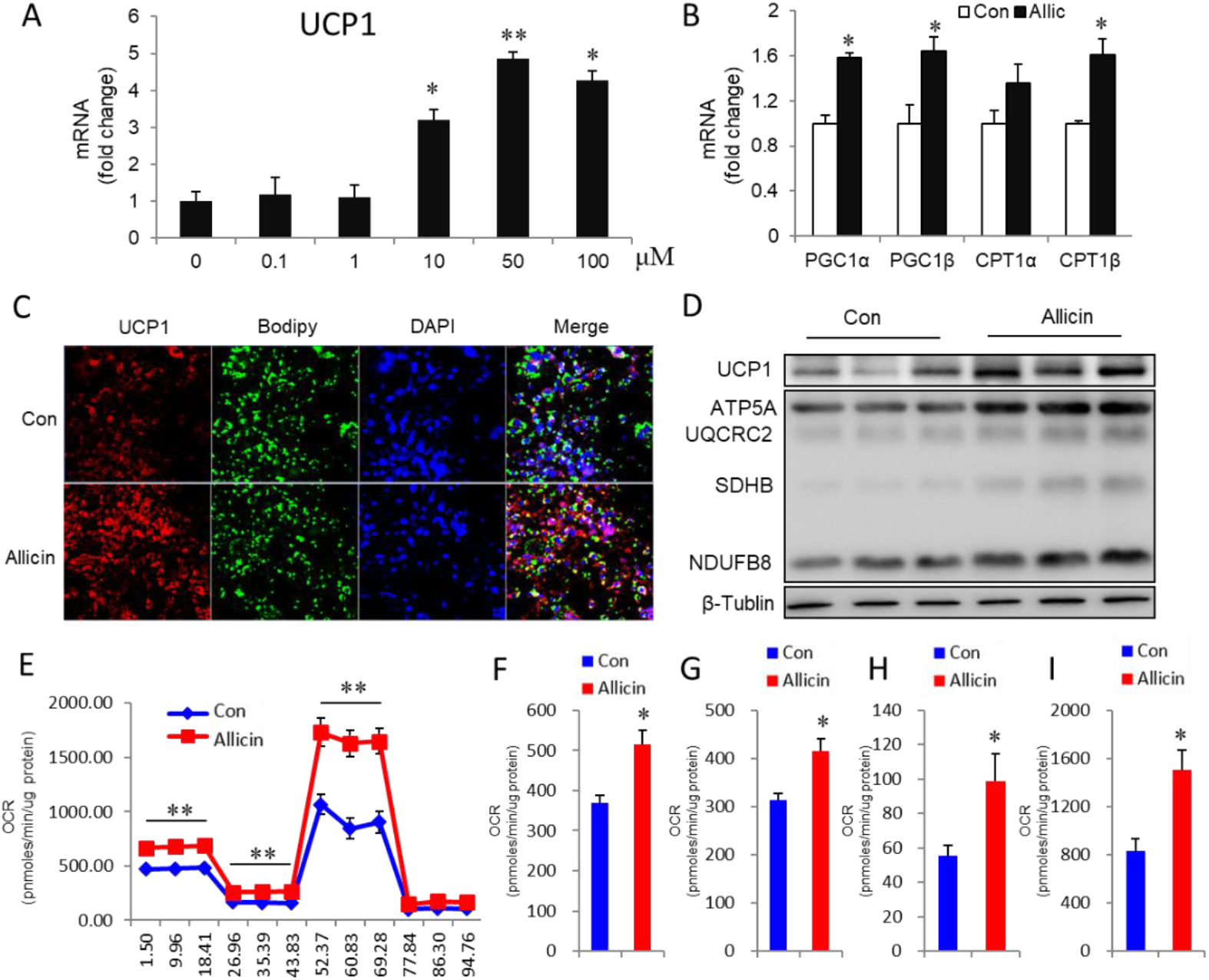
Allicin increases the activity of brown adipocytes and oxygen consumption in vitro. (A) Dose-dependent effect of Allicin on UCP1 expression in primary brown adipocytes at Day 6 of brown adipogenesis. (B) Thermogenic gene expression in brown adipocytes treated with 50 μM Allicin or DMSO. (C) Immunofluorescence staining of UCP1 and Bodipy in brown adipocytes treated with 50 μM rutin or DMSO at day 6 of brown adipogenesis. (D) The protein levels of UCP1 and OXPHOS in brown adipocytes treated with DMSO or Allicin. (E) oxygen consumption rates (OCR) at day 6 of brown adipogenesis with Allicin or DMSO treatment. (n=5). (F-I) The OCR-related basal metabolic rate, ATP levels, maximum oxygen consumption and proton leakage. (n=5). Values represent means ± SEM. Error bars represent SEM; significant differences compared to vehicle controls are indicated by *p < 0.05, **p < 0.01, ***p <0.001 (assessed by Student’s t-test).

## Discussion

Metabolic diseases caused by the unbalance of energy homeostasis are an explosive epidemic, especially obesity, and related metabolic diseases (*27*). BAT is a critical organ that regulates energy homeostasis and known as an important potential target for the treatment of metabolic diseases. Bioactive compounds in food have attracted much attention as a safe and effective molecular library. It has reported that garlic is an excellent natural source of bioactive compounds(*28*). In this study, we revealed that the beneficial effects of Allicin on energy homeostasis in obese mice. Our data show that allicin enhances the BAT function, promotes the energy expenditure, inhibits the fat mass accumulation, improves the glucose homeostasis and insulin sensitivity, and improve hepatic steatosis. Mechanisms, we found that Allicin can induce the promote the activity of sirt1 and induce the downstream PGC1α-Tfam signaling pathway. It is the first study that relatively systematically demonstrated that Allicin induces BAT activation and regulate energy homeostasis. Our findings point out that reasonable proper administration of Allicin may be a promising potential therapeutic strategy for obesity and related metabolic syndromes.

Allicin (thio-2-propene-1-sulfinic acid S-allyl ester) is the principal component of garlic and has many physiological functions. Several studies have reported that garlic or its main ingredient, allicin, has an anti-obesity, anti-hyperinsulinemic, anti-hyperlipidemic, and anti-hypertensive effect (*23, 29–31*). Among them, a research article reported that Allicin could induce brown-like adipogenesis and increased lipid oxidation in subcutaneous fat though KLF15 signal cascade (*23*). However, the effect and mechanism of Allicin on the BAT functions has remained unclear. Additionally, the research of Allicin on regulating energy homeostasis is not systematic. In the present study, our findings demonstrated that Allicin induced BAT activation, leading to the inhibition of obesity and maintain energy homeostasis in mice.

The malfunctions of energy homeostasis, including energy expenditure (EE), food intake, and fat hypertrophy, can cause obesity (*1, 2*). In the current study, Allicin treatment profoundly increased the energy expenditure, reduced fat mass, body weight gain, and improved the glucose homeostasis in HFD mice. Importantly, the similar effect of allicin was further confirmed in DB/DB mice, which is a leptin receptor-deficient genetic obese mice model. This indicates that Allicin has a stable function of preventing and treating obesity and regulating energy metabolism of the body. Furthermore, we investigated the effect of Allicin on BAT activation both in HFD and DB/DB mice and found that Allicin treatment more profoundly promotes the activation of BAT in Db/Db mice than in HFD mice (Fig. 3 and Fig. 4), based on that Db/Db mice are known to deficient in BAT activity(*32, 33*). This result indicated that Allicin has excellent ability to activate BAT and regulate energy metabolism. However, there are many factors (including thyroid hormones, catecholamine neurotransmission, and cytokines) secreted by the endocrine system, which can activate brown adipocyte. So, in order to determine the effect of allicin on BAT activation is direct or secondary, we performed the Allicin treatment on brown adipocytes in vitro. Our data of Allicin treatment in vitro confirmed that Allicin could directly activate the brown adipocyte.

In summary, the present study demonstrated that allicin regulates energy homeostasis through promoting BAT activation. In general, moderate application of Allicin represents a promising potential treatment strategy for prevention of obesity and related metabolic disorder.

## Compliance with ethical standards

The authors declare that they have no conflict of interest.

## Supplementary figure legends

**Figure s1.**
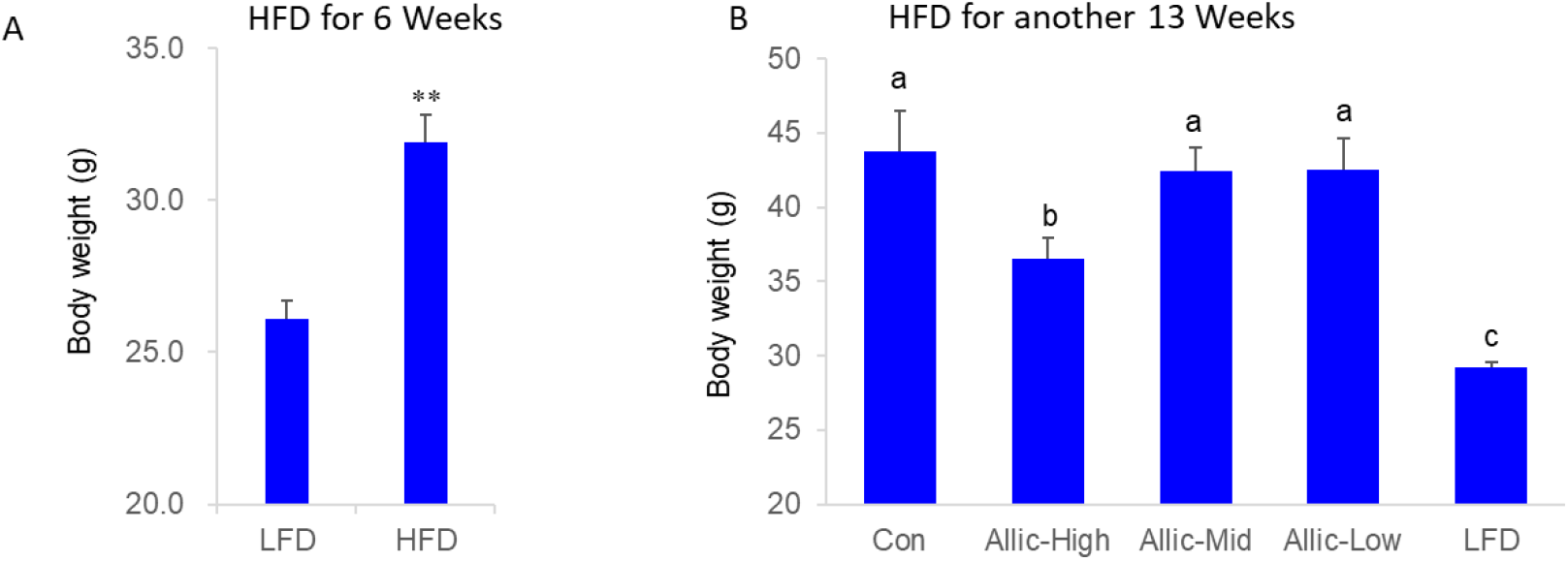
(A) The bodyweight of chow diet and high-fat diet mice for the first 6 weeks before the Allicin treatment. (B)The bodyweight of different group mice with chow diet and high-fat diet for 13 weeks.

**Figure s2.**
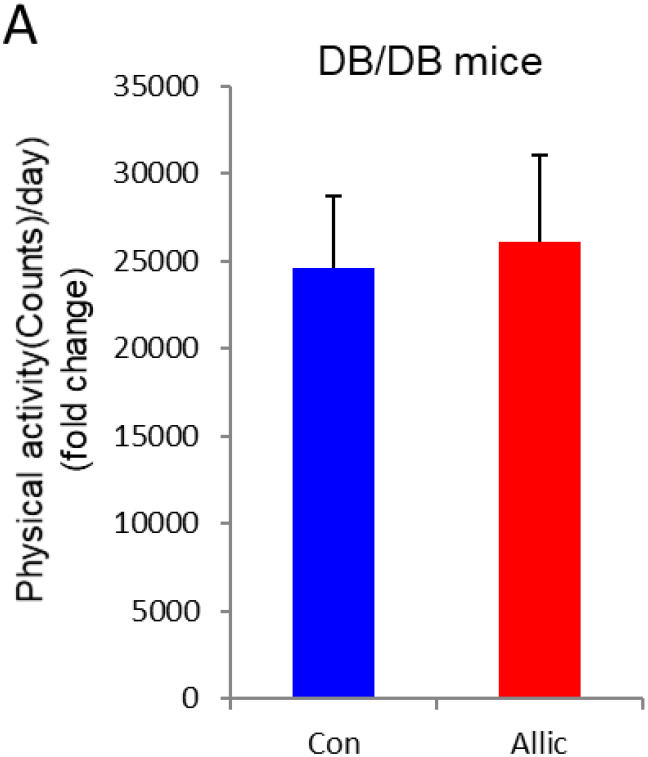
The physical activity of Db/Db mice treated with or without Allicin

**Figure s4.**
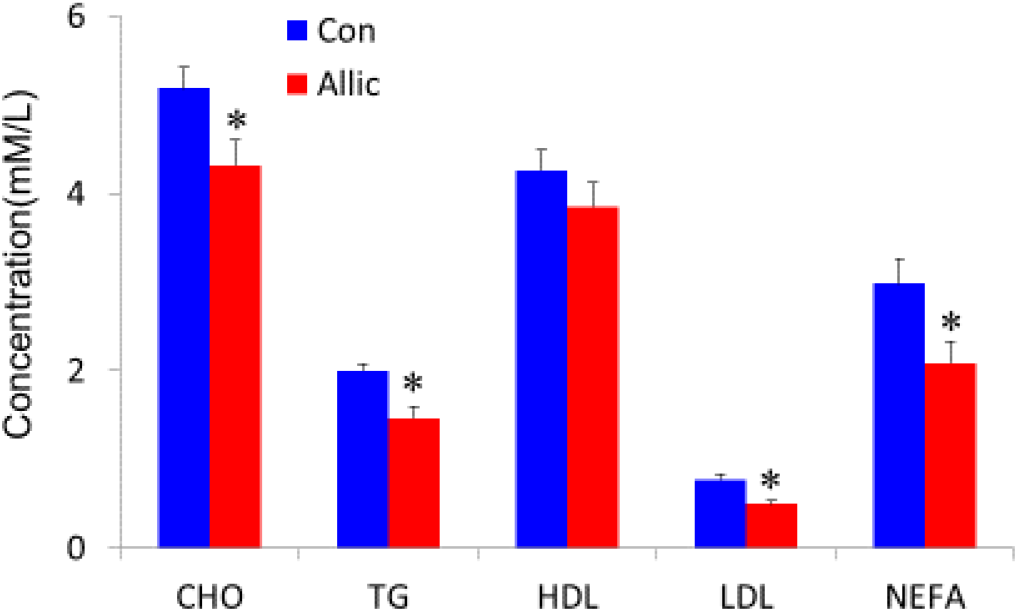
The lipid profile in the serum from control and Allicin-treated DIO mice.

